# Neural representations of task context and temporal order during action sequence execution

**DOI:** 10.1101/2020.09.10.290965

**Authors:** Danesh Shahnazian, Mehdi Senoussi, Ruth M. Krebs, Tom Verguts, Clay B. Holroyd

**Author notes:** equal contribution (Danesh Shahnazian, Mehdi Senoussi).

## Abstract

Since routine action sequences can share a great deal of similarity in terms of their stimulus response mappings, their correct execution relies crucially on the ability to preserve contextual and temporal information (Lashley, 1951). However, there are few empirical studies on the neural mechanism and the brain areas maintaining such information. To address this gap in the literature, we recently recorded the blood-oxygen level dependent (BOLD) response in a newly developed coffee-tea making task (Holroyd et al., 2018). The task involves the execution of 4 action sequences that each feature 6 decision states. Here we report a reanalysis of this dataset using a data-driven approach, namely multivariate pattern analysis (MVPA), that examines context-dependent neural activity across several predefined regions of interest. Results highlight involvement of the inferior-temporal gyrus and lateral prefrontal cortex in maintaining temporal and contextual information for the execution of hierarchically-organized action sequences. Furthermore, temporal information seems to be more strongly encoded in areas over the left hemisphere.

## Introduction

Because many routine action sequences have similar stimulus-response mappings, their correct execution relies on the ability to maintain contextual and temporal information that disambiguates one sequence from another (Lashley, 1951). For example, the sequences for preparing soy milk latte and regular latte are very similar. Like many real-world sequences, they exhibit hierarchical structure: Both are specific instances of making coffee, which itself is an instance of making a hot beverage. A barista who makes both of these drinks regularly must therefore maintain *contextual information* about the identity (soy milk vs. regular milk) of the drink that is currently being made, or else they could confuse the two recipes. Similarly, a common pizza recipe calls for first brushing the dough with olive oil and then spreading the toppings. Therefore pizza-making requires maintenance and manipulation of *temporal information*, i.e., the serial rank order of the actions that have been performed already and the ones that need to be performed later. Notably, the failure to maintain contextual and temporal information can lead to “action slips” (Norman, 1981; Reason, 1990), which are errors that relate to the execution of an action that is inappropriate in the current context but appropriate in the context of other, typically habitual sequences (see Mylopoulos, this issue). Human-like computational solutions to this problem have also been proposed for robotic systems, which confront similar sequencing issues when operating in natural environments (Finzi and Caccavale, this issue).

Despite the importance of temporal and contextual information to everyday behavior, the neural underpinnings and mechanisms that support their maintenance in the service of sequence execution are still debated. While many empirical studies have attempted to address these questions (for a review, see Desrochers & McKim, 2019 and the discussion below), most of these do not feature voluntary production of hierarchical and time-extended sequences, which is said to be essential to complex human behavior (Holroyd and Yeung, 2012). Thus, the ecological validity of their findings is rather limited.

Two recent studies have addressed this gap by using a hierarchically structured temporally extended sequence production task. Balaguer et al., (2016) showed that the anterior midcingulate cortex (aMCC) maintains contextual information in the context of a hierarchical planning task. And in a study that is central to our present investigation, we also found the aMCC to be involved in sequencing (Holroyd et al., 2018). In that study, we recorded the BOLD response in a newly developed coffee-tea making task that required participants to maintain contextual and temporal information during sequence production. The task involves the execution of 4 action sequences that in some steps afford identical responses to identical stimuli. In addition, for each sequence those actions were performed twice over the course of the sequence (i.e. the 3rd and the 5th steps in the sequence, which are labelled as the “stir” actions in each sequence; see Figure. 1A, yellow frames). We tested the prediction, motivated by our previous theoretical work (Holroyd, & Yeung, 2012; Shahnazian & Holroyd, 2018), that aMCC is sensitive to both contextual and temporal information in a hierarchically-organized sequence production task. Toward this end, we trained a recurrent neural network model (Shahnazian & Holroyd, 2018; Botvinick & Plaut, 2004) to perform the task, and we determined the degree of dissimilarity between the patterns of activity across the hidden units of the trained model, for all pairwise combinations of steps and sequences in the task. Application of representational similarity analysis (RSA) (Kriegeskorte et al., 2006, 2008; Freund et al., 2020) confirmed that, consistent with the predictions of the model, the distributed pattern of activity in aMCC codes for the identity of the sequence currently being performed (contextual information), and for the temporal progression through each sequence (temporal information). Moreover, in keeping with past literature (e.g. Desrochers et al., 2019a), an exploratory analysis found that a broad region in rostrolateral prefrontal cortex (RLPFC) codes for contextual information, namely, each of the coffee and tea sequences.

**Figure 1.**
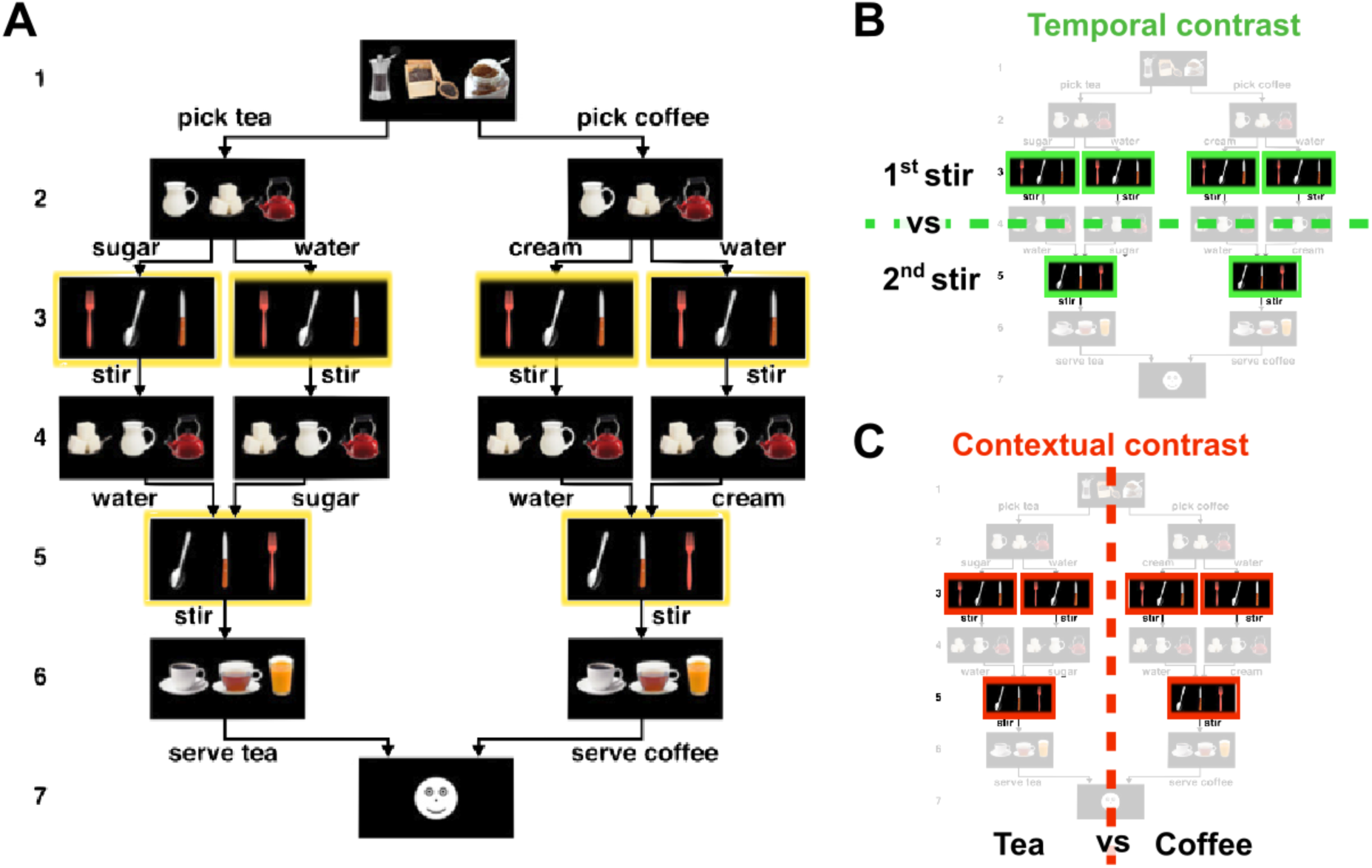
Trial sequence from Holroyd et al. (2018) and contrasts of interest. **A**. The coffee– tea making task. Each trial consisted of a sequence of six decisions followed by feedback (indicated by digits on Left). Each decision comprised three images of different items corresponding to three possible actions (for example, state 1: coffee, pepper, tea; state 2 and 4: cream, sugar, water; state 3 and 5: fork, spoon, knife). For each decision, participants selected one of the three actions by pressing a spatially corresponding button. The position of each object in each picture was randomized across trials. For example, state 1 always entailed a choice between coffee, tea, and pepper, but the locations of these three items varied from trial to trial. Application of the task rules resulted in execution of four different task sequences. Arrows indicate allowable actions. The yellow frames represent the events of interest in the current study. The caption is adapted and summarized from Holroyd et al. (2018). **B**. Temporal contrast: the betas coefficients associated with the first instance of the stirring action event (Class 1) are differentiated from the beta coefficients associated with the second instance of stirring action event (Class 2). **C**. Task contrast: the beta coefficients associated with the stirring action executed in the context of coffee-making (Class 1) are differentiated from the beta coefficients associated with the stirring action executed in the context of tea-making (Class 2).

However, as an hypothesis-based study about the function aMCC, the approach by Holroyd et al. (2018) may have overlooked other brain areas that also code for contextual and temporal information. In particular, the RSA approach adopted by that study is relatively conservative (see discussion). Therefore, here we applied a data-driven approach in order to explore the contribution of other brain areas toward representing contextual and temporal information.

To be specific, we re-analyzed the data from Holroyd et al. (2018) using a different multivariate pattern analysis (MVPA) technique, namely, classification (Haxby et al., 2014), in order to identify the brain regions that specifically represent 1) contextual and 2) temporal information. We classified each dimension of interest (i.e., contextual and temporal information) by applying MVPA on the patterns of activity associated with stir actions, which correspond to steps of the sequence that afford identical responses to identical stimuli (see above). In particular, the stir action states tightly control for potential sensory and motor confounds (Holroyd et al., 2018).

To study the neural topography of contextual and temporal information representations we adopted a region of interest (ROI) approach (Saxe et al., 2006; Poldrack, 2007). This approach consists in first defining functional or anatomical ROIs based on subject-specific data (e.g. gyral-based cortical parcellation) and then performing the analyses of interest, in our case MVPA, in each ROI independently. This ROI approach substantially increases statistical power relative to whole-brain analyses while preserving subject-specific differences in brain organization (see also [Dugué et al., 2017] and [Senoussi et al., 2016] for examples of this approach). We accordingly determined a set of ROIs based on past literature on related questions about sequencing tasks in both humans and nonhuman primates (see discussion). We then extracted anatomical locales for those regions by comparing the coordinates reported in the articles against their approximately corresponding regions, as specified by the automatic gyral-based parcellation algorithm of the brain imaging software package Freesurfer (Desikan et al., 2006). Regarding temporal information, the candidate areas included caudal anterior cingulate cortex and rostral middle frontal cortex (Holroyd et al., 2018; Hyman et al., 2012), caudal middle frontal cortex (Ninokura et al., 2004; Warden & Miller, 2010) middle temporal cortex and pars orbitalis (Kalm & Norris, 2017a), hippocampus (Davachi & DuBrow, 2015), and parahippocampal cortex (Hsieh et al., 2014). Regarding contextual information, the candidate areas included caudal anterior cingulate cortex (Balaguer et al., 2018; Holroyd et al., 2018), caudal middle frontal cortex (Averbeck & Lee, 2007), rostral middle frontal cortex (Desrochers et al., 2019a), hippocampus (Ranganath & Ritchey, 2012), frontal pole (Mansouri et al., 2017) and areas in inferior frontal junction (Wilson et al., 2017) including pars opercularis, pars orbitalis and pars triangularis. We discuss the choice of these areas in the discussion section.

The results provide insight into which brain regions support the execution of action sequences by maintaining contextual and temporal information that disambiguate otherwise identical environmental states or action affordances.

## Methods

A complete description of the participant demographics, materials, experimental procedure and MRI recording can be found in Holroyd et al. (2018). Here we provide a short summary of the relevant aspects.

### Dataset

#### Participants

The experimental procedures were approved by both the Ethical Committee of Ghent University Hospital and the University of Victoria Human Research Ethics Board and the experiment was conducted in accordance with the 1964 Declaration of Helsinki. 18 volunteers (13 female, 5 males) took part in the experiment and provided informed consent. All participants received monetary compensation for their participation.

#### Task

The coffee-tea making task involved production of sequences of six decisions (steps). Each step was prompted by the presentation of 3 images of items arranged in a horizontal row on the computer screen. Participants indicated their response by pressing a spatially corresponding button on the response pad. For each step, a single set of 3 items appeared on the screen, but the spatial locations of the items were randomized, thereby preventing sensory and motor confounds. As a cover story, participants were told that they were baristas, and that on each trial they should choose what beverage (coffee or tea) they would serve to a customer. They were further instructed that to prepare the beverage, two ingredients were to be added. The beverages differed in one ingredient (coffee required milk and tea required sugar) and were identical for the other ingredient (both coffee and tea required water).To prepare the beverage (Figure 1), the participants were instructed to select the drink type (1st decision), add one ingredient (2nd decision), stir the added ingredient (3rd decision), add the other ingredient (4th decision), stir the added ingredient (5th decision), and serve the prepared beverage (6th decision). Participants were instructed to select the beverages and the orders of the ingredients (water or the other ingredient first) in a random manner “as if flipping a coin”. Of particular interest here, the 3rd and 5th decisions, referred to as “stir actions”, were prompted by the presentation of cutlery-related images. Participants on these steps were instructed to always choose the spoon to stir the previously added ingredient, irrespective of the sequence that was being performed. Therefore, the contextual and temporal information cannot be inferred from the sensory input and motor output in these decision states.

#### MRI recording

Structural and functional images were acquired at Ghent University Hospital using a Siemens 3T Magnetom Trio MRI scanner.

#### Preprocessing and Generalized Linear Model

EPI images were 1) aligned to the first image in each time session, 2) corrected for slice acquisition timing; then the mean functional image was co-registered to the individual anatomical volume. The co-registered images were not normalized or smoothed.

A general linear model was then applied to the resulting voxel-level time-series in each block. The regressors of this analysis were generated through convolving the time-series impulse function for each of the 24 task states (6 decisions across 4 sequences) as well as motion parameters and global signal, with a canonical hemodynamic response function.

### Multivariate pattern analysis

#### Contrasts of interest

As will be detailed below, we used a linear support vector machine (SVM) classifier, implemented in the Scikit-learn Python toolbox version 0.23.1 (Pedregosa et al., 2011), to classify activity patterns associated with each event of interest in each of the predetermined ROIs (De Martino et al., 2008).

In the temporal contrast, we trained the SVM classifier on the normalized beta coefficients (Kalm & Norris, 2017b) associated with actions 3 and 5 (i.e. stir actions) in the sequence, irrespective of the performed sequence (i.e. coffee or tea sequence), to discriminate between the first and second instance that the stir action was performed. For the context contrast, we trained a classifier on the same beta coefficients to discriminate whether the stir action was performed in a tea or a coffee sequence, irrespective of the sequence position of the stir action (i.e. first or second stir). Given that the GLM provides beta estimates for 2 consecutive stir actions across 4 sequences in each of the 4 blocks, there were a total of 32 observations (16 per class) to be classified for each participant in either of the contrasts (Figure 1B-C).

#### Controlling for univariate confounds

A recent study by Kalm & Norris (2017b) has shown that in studies investigating the neural correlates of sequence learning or sequence execution, an important confounding factor may be the concurrent evolution of other cognitive factors such as cognitive load or general drift in neural activity throughout a sequence. They argue that effects can mask actual sequence representations and confound results. To control for such confounding factors, we followed one of their suggestions and normalized beta values associated with each event of interest, independently for each participant and ROI. More specifically, in each classification analysis, for each of the 32 event-related betas to be classified, we subtracted the average beta value of all voxels in that ROI in this event, and divided by their standard deviation.

#### Regions of interest

As discussed above, our analysis focused on a set of predetermined ROIs as defined by the Freesurfer automatic gyral parcellation algorithm (Fischl, 2012; Desikan et al., 2006). For each of these ROIs only voxels located in the gray matter, as segmented by the automatic gray-white matter segmentation by Freesurfer, were used for analyses. The ROIs include areas in lateral frontal cortex and frontal operculum (frontal pole, pars opercularis, pars orbitalis and pars triangularis, rostral and caudal middle frontal cortex); in medial prefrontal cortex (caudal anterior cingulate cortex); and in the temporal lobe (middle temporal cortex, hippocampus and parahippocampal cortex, see Figure S1 in supplementary material). To enable comparisons across contrasts, all of these ROIs were subjected to the same set of analyses.

#### Regions of interest analysis

For the classification analysis in each ROI, we carried out a feature-selection procedure in which we first conducted a univariate analysis of variance of all the voxels with the contrast of interest as the independent variable and the beta-values associated with the training set’s observations as the dependent variable, and then selected the 120 voxels (see Figure S2 which illustrates the effect the number of features has on the classifier’s performance) with the highest F-statistics as features for the classification analysis (Zhang et al., 2013). We then fit the linear SVM to classify the beta-values and took the average 4-fold cross validated classification accuracy as the accuracy score for that ROI. The feature-selection and classification were performed in a cross-validation procedure to avoid double-dipping, using the *cross_validate* and *Pipeline* functions from Scikit-learn.

We additionally carried out a second ROI classification analysis to test whether temporal and contextual neural representations generalize across different tasks (coffee vs. tea) and step in the sequence (3rd vs 5th) respectively. To do so we modified the cross-validation scheme for each contrast as follows: In the temporal contrast, the classifiers were trained to differentiate the first and second stir actions of coffee sequences, and subsequently tested on first and second stirs of tea sequences for the first cross-validation fold, and the reverse for the second cross-validation fold. For the contextual contrast, classifiers were trained to differentiate coffee and tea sequences in the first stir action-related betas in the first two blocks of the session, and tested on the second stir action-related betas in the last two blocks of the session, and vice versa. We drew the training and test data for the context contrast from different experimental blocks in order to control for slow BOLD changes that could confound the analysis. These analyses allowed us to test whether the discriminative multivariate neural patterns representing the temporal and contextual information generalize across the sequence tasks and steps, respectively (see Bernardi et al., 2018).

To assess the significance of the resulting classification accuracies we carried out a one-sided binomial test on the scores for each ROI separately (across participants). The binomial test assesses the probability of obtaining the observed number of “successes”---correct classification of a pattern at test in our case---based on the probability of a success in one trial at chance-level performance (50%) and the number of “trials”, i.e. the total number of classifier predictions. Given that for each of the 18 participants 32 betas were classified at test, there were a total of 576 “trials” in our second-level (i.e. group-level) analysis. Previous research (Combrisson & Jerbi, 2015) has shown that the binomial test is a better estimator of classification accuracy significance and is a more appropriate statistical method than the Student T-test because classification accuracies do not follow a normal distribution (i.e. they are 0 and 1). A false discovery rate (FDR) correction (Benjamini & Hochberg, 1995) was then used to correct for multiple comparisons across ROIs (10 multiple comparisons), for each contrast.

Furthermore, we carried out an exploratory analysis on hemispheric differences in contextual and temporal information representations. We performed the same ROI classification analysis on all of the ROIs except for the frontal pole ROI (as 8 participants had 30 or fewer voxels in either the left or right hemisphere of that ROI). Thus, on the 9 remaining ROIs we carried the ROI classification analysis considering each hemisphere separately. To assess the statistical significance of these results, ROI-by-hemisphere classification accuracies were entered into a 2×2×9 repeated measures Anova with factors contrast (2 levels: temporal, contextual), hemisphere (2 levels: left, right) and ROI (9 levels: parahippocampal, hippocampus, caudal anterior cingulate, middle temporal, caudal middle frontal, rostral middle frontal, pars opercularis, pars triangularis, pars orbitalis).

### Data and code availability

All raw and preprocessed data are available from the Open Science Framework repository created by Holroyd et al. (2018): https://osf.io/wxhta/. All analysis scripts created for the current study are available on the Github repository: https://github.com/mehdisenoussi/EST.

## Results

Participants successfully implemented the task instructions (see Holroyd et al., 2018). No trials were rejected, so our current analysis is not biased because of differences in error patterns.

### ROI classification

For the temporal contrast, the analysis of cross-validated classification scores (Figure 2) revealed significantly higher than chance-level performance (after FDR correction), in the following ROIs: parahippocampal cortex (median ± [1st quartile, 3rd quartile]; 53.65% ± [46.87, 56.25]), middle temporal cortex (60.94% ± [57.03, 64.84]), caudal (62.50% ± [53.91, 68.75]) and rostral (65.62% ± [57.03, 68.75]) middle frontal cortex, pars opercularis (56.25% ± [50.78, 59.37]), pars triangularis (62.50% ± [53.91, 68.75]), and pars orbitalis (57.81% ± [53.12, 62.5]). In the context contrast, classification analysis (Figure 2) revealed significantly higher than chance-level performance (after FDR correction), in pars triangularis (53.12% ± [45.31, 59.37]) and pars orbitalis (57.81% ± [46.87, 71.09]). (In our explorations, we examined the ROIs using a searchlight classification approach [Senoussi et al., 2016], results are largely the same [Figure S3.]). (Note that for some participants, not all ROIs contained 120 voxels or more, for these ROIs, the classification analysis was performed with all the voxels available, see Figure S4.)

**Figure 2.**
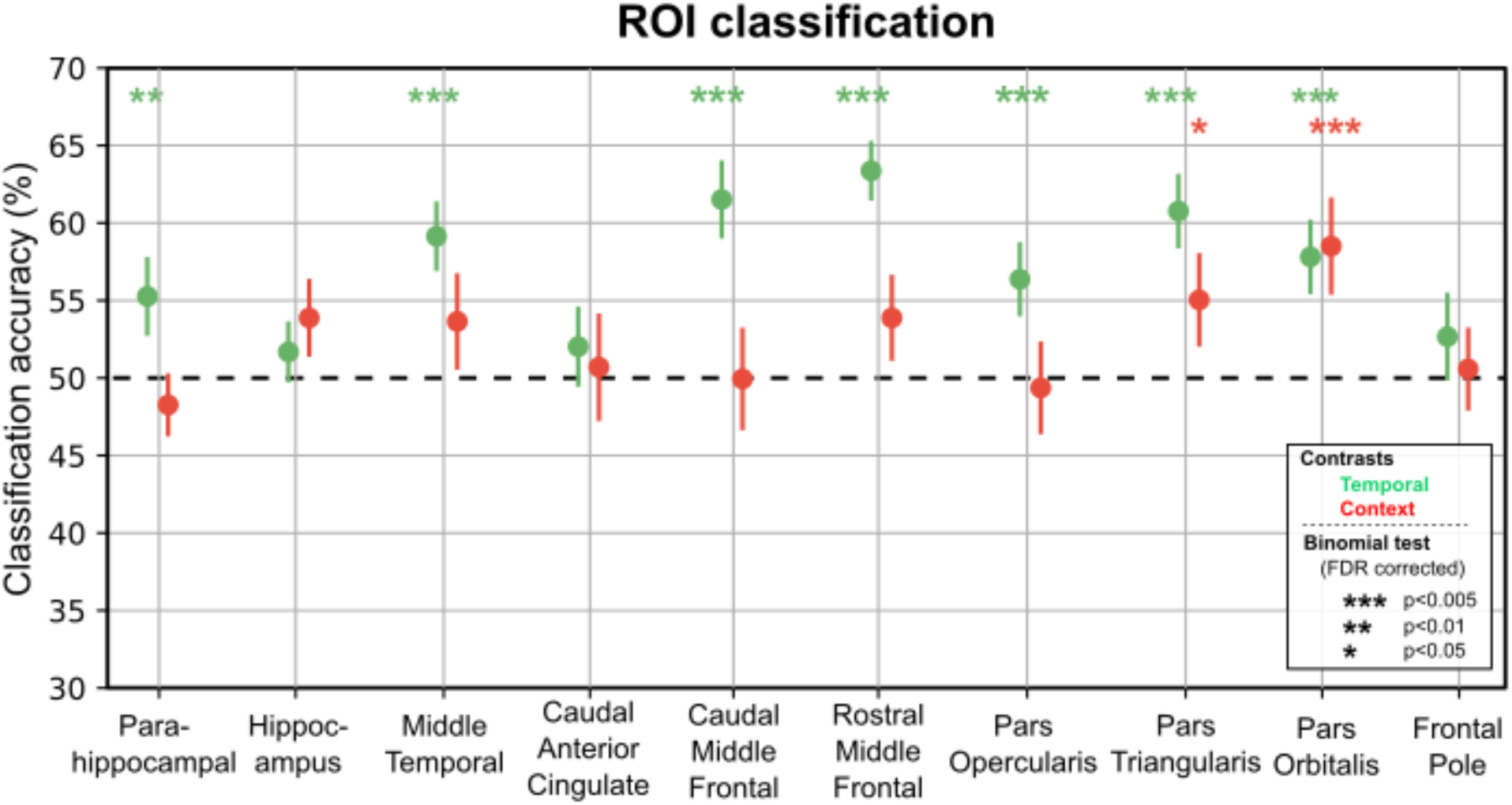
ROI classification accuracy. Results of the classification accuracy for each ROI using feature selection (k=120 voxels), by contrast. Colors represent contrasts (green for temporal contrast, red for context contrast). Significance is represented by color-coded stars above each contrast/ROI (see legend inside the graph).

### Analysis of the generalizability of the discriminative patterns

The results of the generalizability analysis (Figure 3, see descriptive statistics in the supplementary material) were highly similar to the ROI classification analysis (Figure 2): For the temporal contrast, the generalizability analysis identified most of the ROIs (except for parahippocampal cortex) that were previously found in the ROI classification analysis (i.e. middle temporal cortex, caudal and rostral middle frontal cortex, pars opercularis, pars triangularis and pars orbitalis) (after FDR correction). However, this analysis also identified the hippocampus and caudal anterior cingulate cortex, which were not revealed by the ROI classification analysis. For the context contrast, the generalizability analysis identified both pars triangularis and pars orbitalis (after FDR correction), in keeping with the ROI classification analysis.

**Fig. 3.**
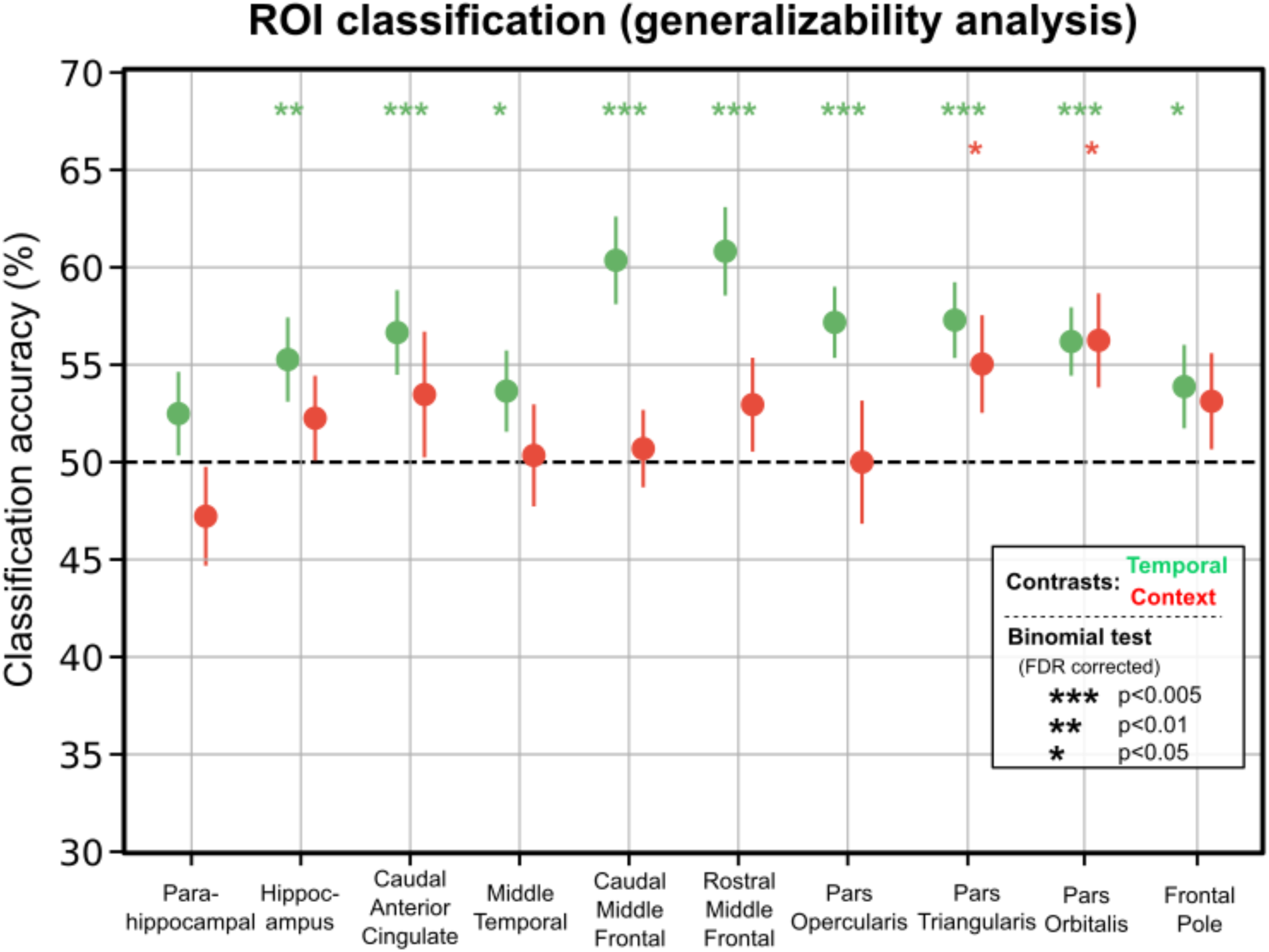
ROI classification accuracy: generalizability of the discriminative patterns. Results of the classification accuracy on per ROI using feature selection (k=120 voxels) in the generalizability analysis. Colors represent contrasts (green for temporal contrast, red for context contrast). Significance is represented by color-coded stars above each contrast/ROI (see legend).

### Hemispheric analysis

The ROI-hemisphere analysis of cross-validated classification accuracies with factors hemisphere, ROI and contrast, revealed a main effect of ROI (*F*(8, 136) = 2.67, *p* = 0.0093) and no main effects of hemisphere or contrast (Figure 4). Additionally, we found significant two-way interaction between contrast and ROI (*F*(8, 136) = 2.65, *p* = 0.0099), as well as two-way interaction between contrast and hemisphere (*F*(1, 17) = 4.87, *p* = 0.0414). To explore what underlies the significant interaction between contrast and hemisphere we investigated the difference between the average classification accuracy for each contrast by hemisphere across ROIs. There was a marginal difference in classification accuracy between the temporal and contextual contrasts in the left hemisphere (with classification accuracies in the temporal contrast being higher than the ones in contextual contrast; *t*(17) = 1.93, *p* = 0.0699), but no consistent difference in the right hemisphere (*t*(17) = 0.49, *p* = 0.6783).

**Figure 4.**
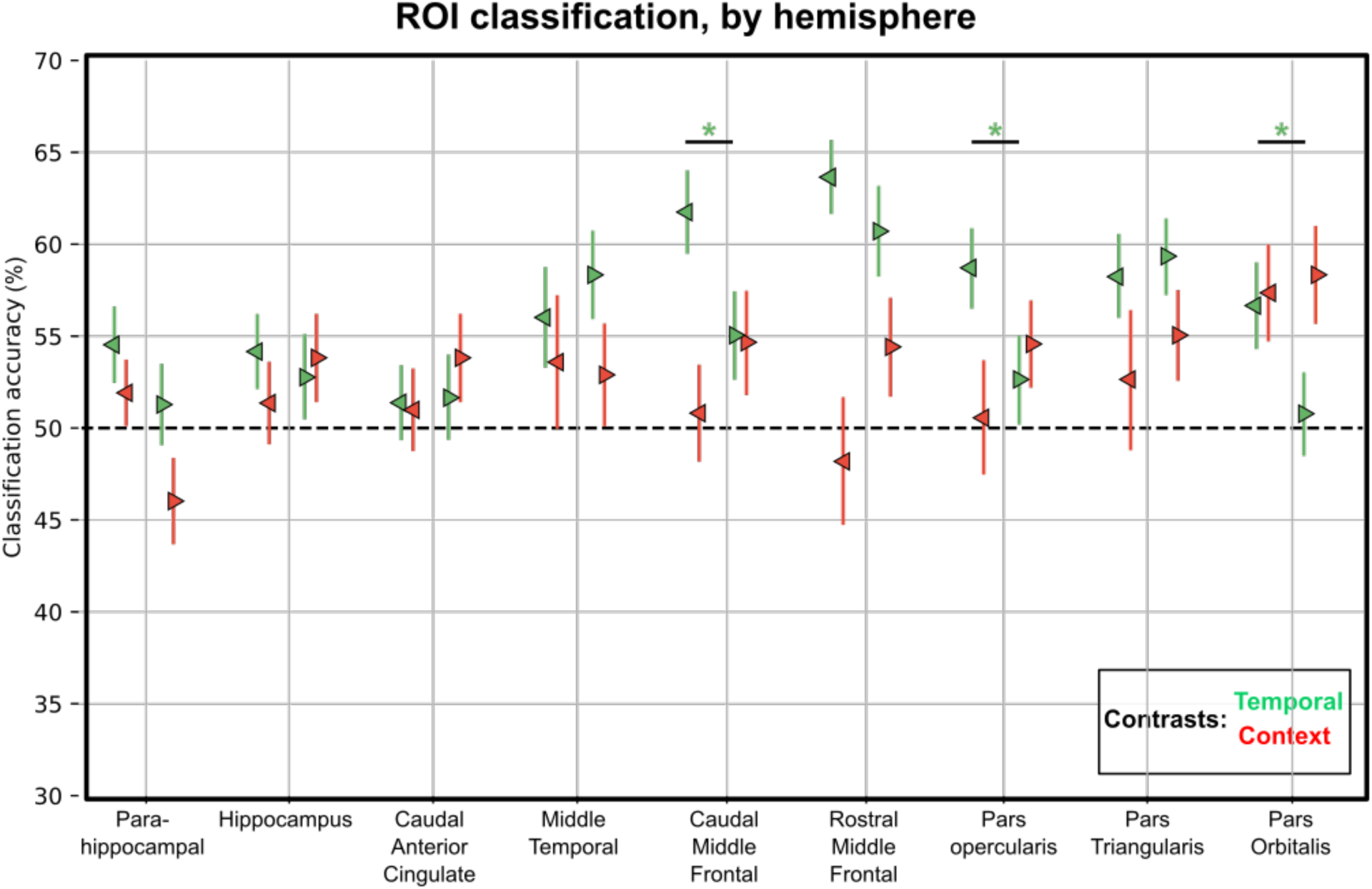
ROI classification accuracy by hemisphere. Left pointing triangles represent ROIs of the left hemisphere, right pointing triangles represent ROI of the right hemisphere. Black bars and stars represent significant differences between classification accuracy of Left and Right hemispheres of an ROI as revealed by the post-hoc tests.

Further post-hoc tests were conducted only on the ROIs that showed at least one significant classification accuracy (either in temporal, or context contrast, or both; *p*<0.05, FDR corrected). For each ROI, we compared the classification accuracies of the left and right hemisphere in each contrast with a two-sided paired samples T-test. For the temporal contrast, this analysis revealed that classification accuracies were higher in the left hemisphere in caudal middle frontal gyrus(*t*(17)=2.83, *p*=0.0115), pars opercularis (*t*(17)=2.5, *p*=0.023) and pars orbitalis (*t*(17)=2.30, *p*=0.0341). All other ROIs in the temporal contrast did not show significant differences between classification accuracies obtained from the left and right hemispheres (all *ps* > 0.25). For the contextual contrast, the only ROI exhibiting significantly above-chance performance in the ROI classification analysis was pars orbitalis and it did not show any significant difference between hemispheres (*p*>0.7). The descriptive statistics associated with the post-hoc analyses can be found in the supplementary material.

## Discussion

In this inquiry, we examined which brain areas code for contextual and temporal information necessary for voluntary sequence production. We used pattern classification to detect brain areas that encode states of temporally extended, hierarchically structured sequences differently depending on which sequence is being performed (task contrast) or which step of each sequence is being implemented (temporal contrast).

The temporal contrast revealed that areas in the IFG (e.g. pars triangularis), lateral prefrontal cortex (caudal and rostral middle frontal areas) and temporal lobe (parahippocampal cortex, middle temporal cortex) code for temporal information in this task.

The finding that lateral prefrontal cortex is involved in representing temporal information during sequence execution are consistent with studies in monkeys showing that the activity of neurons in the prefrontal cortex (PFC) is differentially modulated by the rank order of the actions in a sequence (Ninokura et al., 2004; Warden &Miller, 2010; Carpenter et al., 2018). Similarly, other studies have shown that the activity in rostrolateral and dorsolateral prefrontal cortex (RLPFC) (Desrochers et al., 2019a; Averbeck et al., 2003) ramps up in direct relation to the sequence progression, presumably as a means to encode temporal information. Also consistent with our findings are previous MVPA fMRI studies that have revealed several brain areas, including the anterior temporal lobe, RLPFC and pars orbitalis and parahippocampal cortex (Kalm & Norris, 2017a, Hsieh et al., 2014), that are sensitive to the rank order of sequential events. By contrast, inconsistent with the past literature, in the ROI analysis, we did not find evidence for involvement of the aMCC and hippocampus (although both of these areas are revealed in the generalizability analysis) in the distributed encoding of temporal information (for example, see Procyk et al., 2000; Ma et al., 2014; Davachi & DuBrow, 2015). We also found indications that the temporal information can be more strongly decoded in the left hemisphere, which may suggest some degree of lateralization of this type of representation.

These results were bolstered by the generalizability analysis, which showed that the pattern reflecting time-information in one task (e.g. coffee making) can also be used to discern time information in the other task (e.g. tea making) in most of the areas detected in the ROI analysis. This indicates that the time-information is partly represented in an abstract (task-invariant) fashion. Only two ROIs survived FDR correction for multiple comparisons in the analysis of the context contrast (and also the generalizability analysis), namely pars orbitalis and pars triangularis. This is consistent with some theoretical work that attributes a domain-general role to pars orbitalis in learning the relationship between environmental events and transition probabilities between various environmental states (Wilson et al., 2017; Niv, 2019). Similarly, pars triangularis is shown to be involved in learning artificial grammars---which is a form of sequence learning---and verbal working memory (Uddén & Bahlmann, 2012; Price, 2012). The null findings in the other ROIs are contrary to past studies that found evidence for rostral lateral prefrontal cortex (Averbeck & Lee, 2007; Desrochers et al., 2015), aMCC (Powell & Redish, 2016; Balaguer et al., 2016; Holroyd et al., 2018) and the hippocampus in maintaining contextual information (Gupta et al., 2010; Ranganath & Ritchey, 2012). The null findings also contrast with theoretical work positing that aMCC and frontopolar cortex should be involved in the maintenance of contextual information (for example see: Shahnazian & Holroyd, 2018; Mansouri et al., 2017).

One reason for our null findings may be that the locale and the expanse of different anatomical areas offered by the Freesurfer parcellation does not closely match the localization of different sequencing related functions within the frontal cortex. For example, the delineation of the anatomical regions in lateral frontal cortex according to the Freesurfer nomenclature creates very large ROIs in comparison with the ROIs in IFG or the frontopolar ROI. Although we used feature selection to mitigate this problem, the differences in size and the mismatch between anatomical and functional separation of brain areas may have limited our ability to detect the representation of contextual information in some brain areas.

Notably, we did not find evidence that caudal anterior cingulate cortex codes for contextual information. In the temporal contrast, we were able to detect the caudal anterior cingulate cortex only in the generalizability analysis. This is despite the fact that RSA analysis of the same dataset shows that the aMCC (which overlaps anatomically with the caudal anterior cingulate cortex) is involved in representing contextual information. As mentioned before, one reason might be that the anatomical parcellation of the caudal anterior cingulate cortex does not well match the location and the expanse of the functional area often referred to as aMCC. Another reason may be that RSA and pattern classification approaches have different sensitivity profiles to pattern dissimilarities (Walther et al., 2016).

Perhaps most importantly, the aMCC cluster in the previous study was identified following a combined searchlight/RSA approach, where the predictive RDM was derived from the activation states of the hidden units of an RNN trained on the sequencing task, for all possible states of the task. Because the internal representations of RNNs are typically distributed and exhibit multi-selectivity (Holroyd & Verguts, manuscript submitted), the aMCC cluster reflects these properties. Therefore, even though post-hoc analyses confirmed that this cluster was sensitive to temporal and contextual information, neither of these two factors on their own appear to have sufficient statistical power to yield much aMCC activity. Furthermore, the present study focused on only a subset of task states (whereas the RSA evaluated the pattern of dissimilarity across **all** time-steps; Holroyd et al., 2018) and normalized the data in order to prevent potential confounds that naturally result from multi-selectivity (Kalm & Norris, 2017b). The present study confirms that the aMCC’s distributed code for temporal information is not sufficiently consistent across blocks and sequence types to be detectable using the ROI classification approach, revealing only a weak effect size in the generalization analysis. The two approaches therefore appear to provide complementary information.

Most of the past literature on sequencing (Desrochers & McKim, 2019b) uses tasks that do not emulate the hierarchical and temporally-extended nature of routine behavior. One reason for this is perhaps that sequence execution engages a variety of general domain processes including working memory, action selection, cognitive effort and performance monitoring. Disentangling such general domain processes from the ones that are **uniquely** important to successful implementation of internally guided sequences of actions requires tight control over various confounds, which is hard to achieve when studying hierarchical and extended sequences. These results and those of our previous work (Holroyd et al., 2018) and of Balaguer et al. (2018) are notable exceptions that generalize better to routine behavior in real life. We therefore advocate for adoption of similar tasks in future enquiries on the neural underpinnings of routine behavior. We also suggest that a combination of RSA and pattern classification analysis should be used to examine various aspects of the distributed pattern of activity in various brain areas.

## Acknowledgements

This project was supported in part with funding by the European Research Council (ERC) under the European Union’s Horizon 2020 research and innovation programme (Grant agreement No. 787307 awarded to CBH and 636116 awarded to RMK). MS and TV were supported by grant G012816 from Research Foundation Flanders.

## Supplementary figures

**Figure S1.**
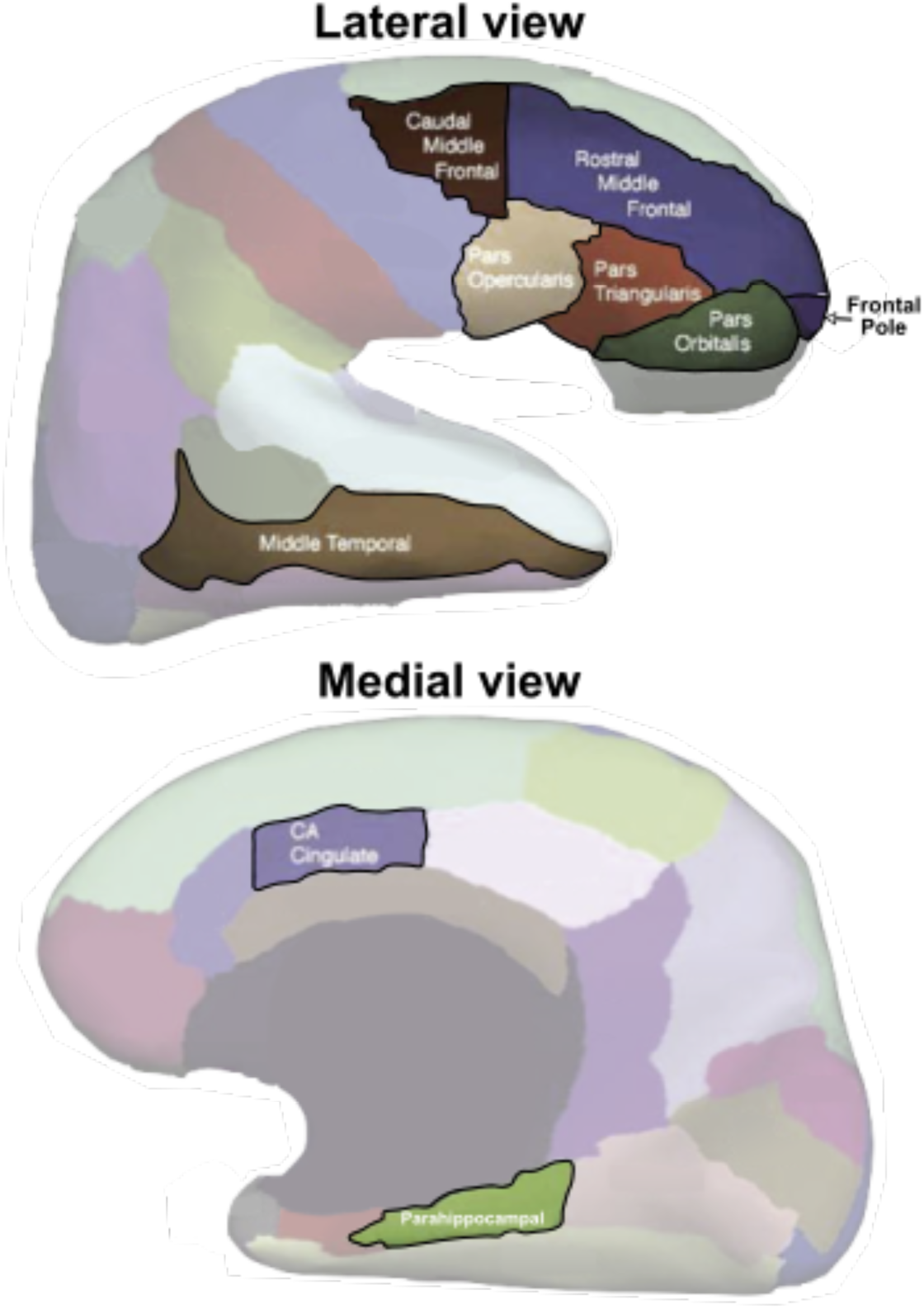
Regions of interest. The anatomical locations of 9 of the 10 ROIs used in the analyses: frontal pole, pars opercularis, pars orbitalis and pars triangularis, rostral and caudal middle frontal cortex, caudal anterior cingulate cortex, middle temporal cortex and parahippocampal cortex. The hippocampus ROI was obtained from another parcellation performed by Freesurfer, it is thus not represented in this figure. The figure represents lateral (top) and medial (bottom) views of an inflated brain. Note that the inflated brain is used for illustration purposes, the ROIs were extracted and used in 3D representation of the data, not on the surface format. (Note that this figure is pending permission from the Elsevier group).

**Figure S2.**
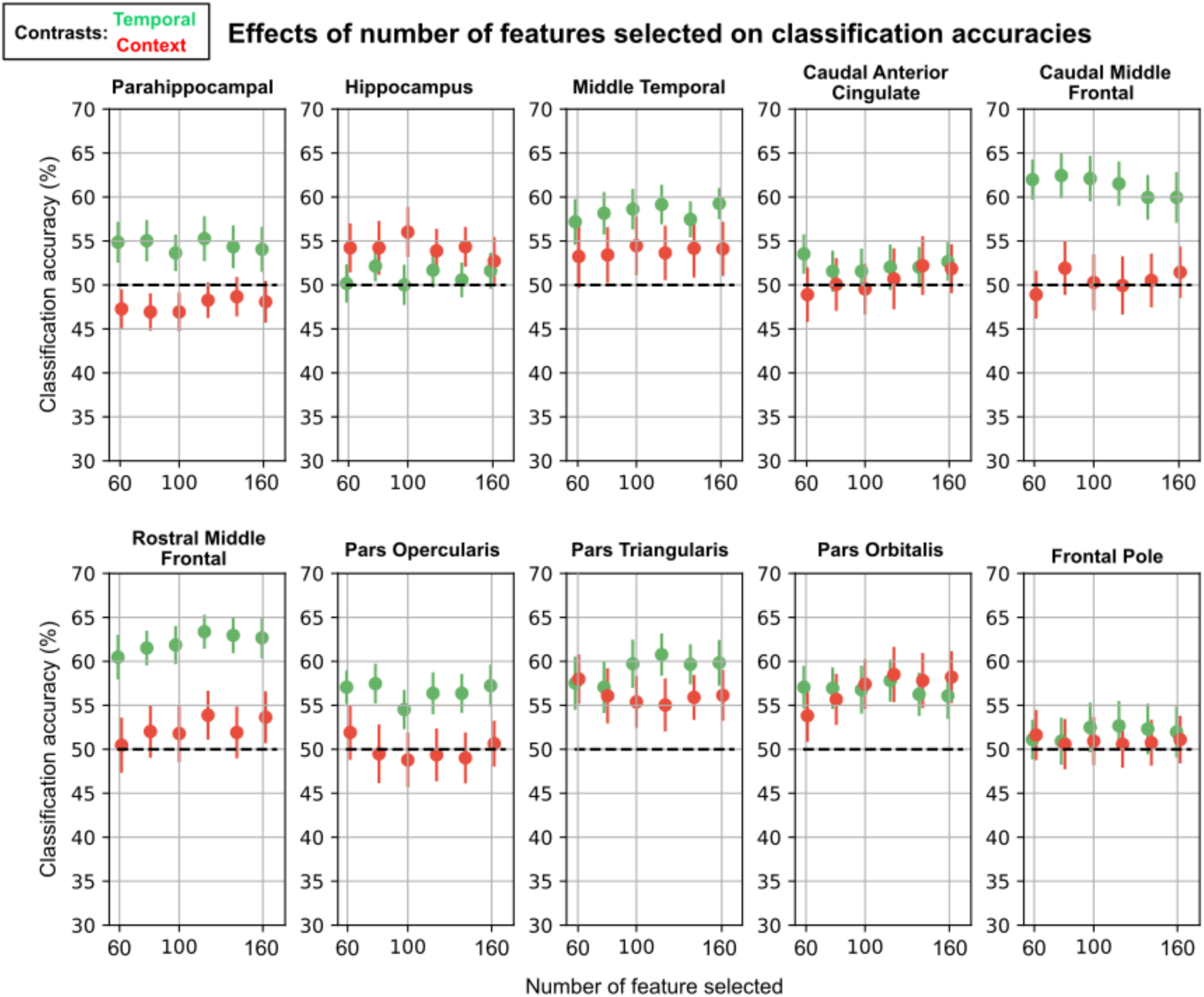
Varying the number of features used for feature selection in ROI classification. We varied the number of features selected for each ROI for classification purposes to verify the stability of classification results across different numbers of selected features. These results reproduce the main findings using 120 voxels for feature selection: it is possible to classify temporal contrast in parahippocampal cortex, middle temporal cortex, caudal middle frontal cortex, rostral middle frontal cortex, pars opercularis, pars triangularis and pars orbitalis; whereas only pars orbitalis and triangularis exhibited stable above-chance classification accuracy for the context contrast.

**Figure S3.**
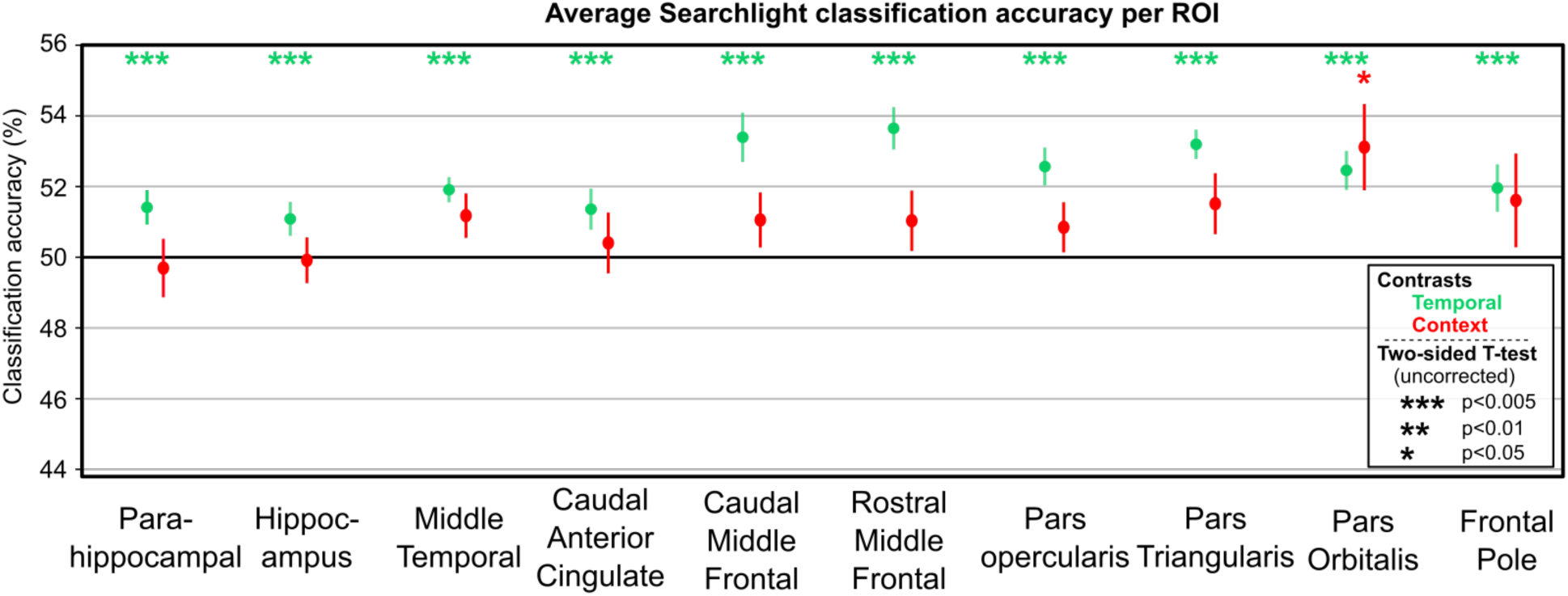
Average searchlight classification accuracy per ROI. Results of the average searchlight classification accuracies across all voxels of each ROI for each contrast. A searchlight of 6mm was used for this analysis. Colors represent contrasts (green for temporal contrast, red for context contrast). Significance was computed using a Student T-test against chance (50%) since the average scores for each ROI are constituted of non-independent samples which violates the binomial test hypothesis.

**Figure S4.**
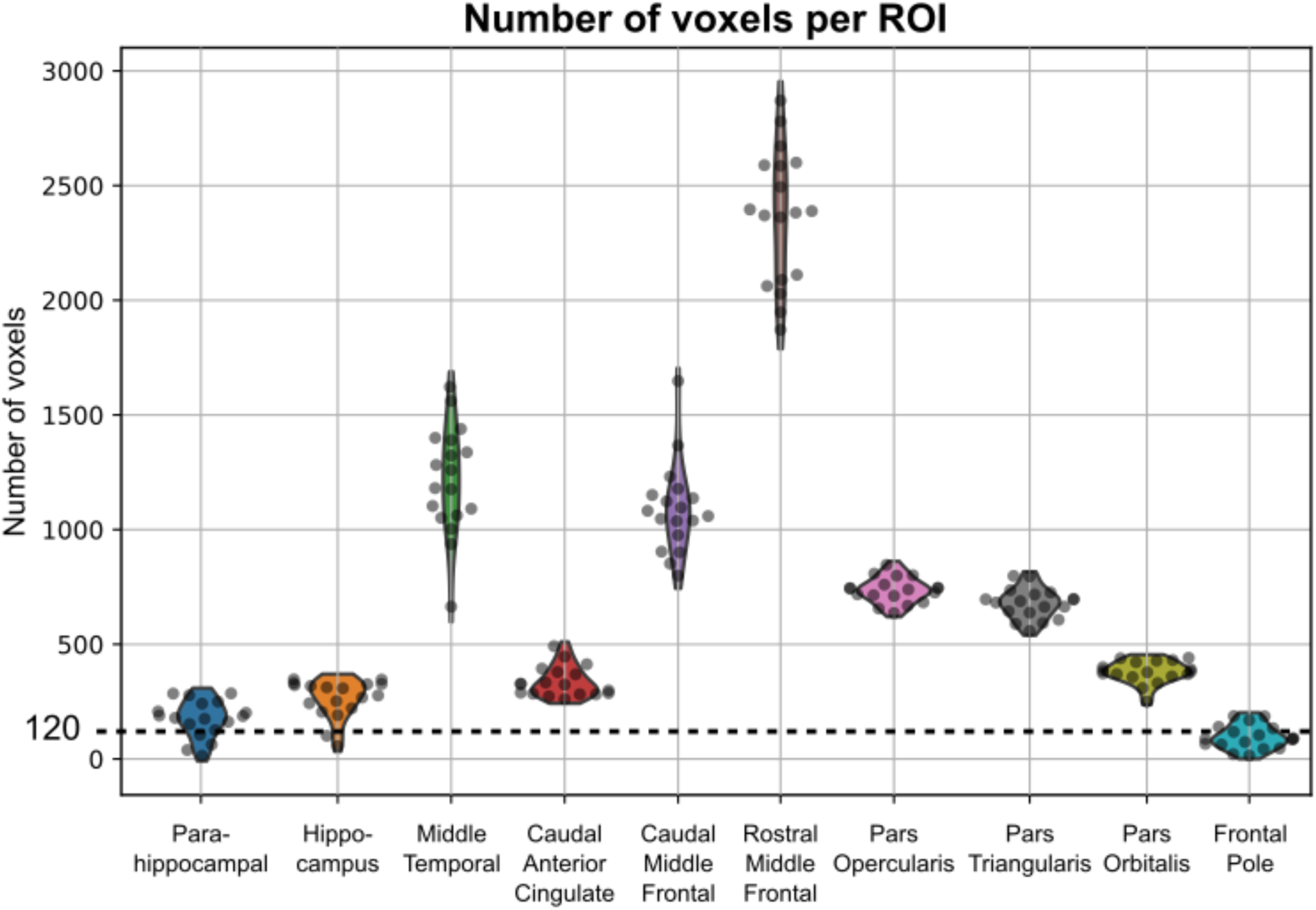
Number of voxels extracted from each ROI per participant. The dashed line represents 120 voxels which is the default number of voxels selected in the feature selection procedure for classification purposes. Each colored shape is a violin plot representing the distribution of number of voxels per ROI across participants computed using a gaussian kernel density estimation. Each dot represents the number of voxels for a participant.

The searchlight analysis of pattern classification (Kriegeskorte et al., 2006) in spherical searchlights of 6 mm radius averaged per ROI (Senoussi et al., 2016) revealed that for the temporal contrast, the averaged classification accuracy of all ROIs were above chance-level performance for the temporal contrast (all p<0.005) and that only classification accuracies of voxels in Pars Orbitalis were above chance-level for the context contrast (p<0.05).

## Post-Hoc tests on the effect of hemisphere on classification accuracy in each ROI

**Temporal contrast** (T-test on related samples: Left versus Right hemisphere):

- <u>parahippocampal cortex</u>: t(17)=1.16, p=0.258
- <u>hippocampus</u>: t(17)=0.53, p=0.601
- <u>caudal anterior cingulate</u>: Neither hemisphere was significantly above chance.
- <u>middle temporal</u>: t(17)=-0.76, p=0.453
- <u>caudal middle frontal</u>: **t(17)=2**.**83, p=0**.**011**
- <u>rostral middle frontal</u>: t(17)=1.13, p=0.283
- <u>pars opercularis</u>: **t(17)=2**.**5, p=0**.**023**
- <u>pars triangularis</u>: t(17)=-0.36, p=0.714

**Contextual contrast** (T-test on related samples: Left versus Right hemisphere):

- <u>pars orbitalis</u>: t(17)=-0.26, p=0.791

### Generalizability analysis descriptive statistics

**Temporal contrast** (median ± [1^st^ quartile, 3^rd^ quartile]

- middle temporal cortex (60.94% ± [48.44, 62.5])
- caudal middle frontal cortex (59.4% ± [56.25, 65.63])
- rostral middle frontal cortex (57.81% ± [54.69, 65.63])
- pars opercularis (56.25% ± [53.13, 62.5])
- pars triangularis (57.81%, ± [50.0, 59.38])
- pars orbitalis (56.25% ± [53.13, 61.72])
- hippocampus (55.21% ± [50.0, 64.06])
- caudal anterior cingulate cortex (56.25% ± [50.0, 59.38])

### Context contrast

- pars triangularis (50.0% ± [50.0, 61.71875])
- pars orbitalis (56.25 % ± [48.44, 64.843])

